# Common neurodegeneration-associated proteins are physiologically expressed by antigen-presenting cells and are interconnected via the inflammation/autophagy-related proteins TRAF6 and SQSTM1

**DOI:** 10.1101/690677

**Authors:** Serge Nataf, Marine Guillen, Laurent Pays

## Abstract

There is circumstantial evidence that, under neurodegenerative conditions, peptides deriving from aggregated or misfolded specific proteins elicit adaptive immune responses. On another hand, several genes involved in familial forms of neurodegenerative diseases were found to exert key innate immune functions. However, whether or not such observations are causally linked remains unknown. To start addressing this issue, we followed a systems biology strategy based on the mining of large proteomics and immunopeptidomics databases. First, we retrieved the expression patterns of common neurodegeneration-associated proteins in two professional antigen-presenting cells, namely B-cells and dendritic cells. Surprisingly, we found that under physiological conditions, numerous neurodegeneration-associated proteins are abundantly expressed by human B-cells. A survey of the human proteome allowed us to map a unique protein-protein interaction network linking common neurodegeneration-associated proteins and their first shell interactors in human B-cells. Surprisingly, network connectivity analysis identified two major hubs that both relate with inflammation and autophagy, namely TRAF6 (TNF Receptor Associated Factor 6) and SQSTM1 (Sequestosome-1). Moreover, the mapped network in B-cells also comprised two other hub proteins involved in both inflammation and autoimmunity: HSPA8 (Heat Shock Protein Family A Member 8 also known as HSC70) and HSP90AA1 (Heat Shock Protein 90 Alpha Family Class A Member 1). Based on these results, we then explored the Immune Epitope Database “IEDB-AR” and actually found that a large share of neurodegeneration-associated proteins were previously reported to provide endogenous MHC class II-binding peptides in human B-cells. Of note, peptides deriving from amyloid beta A4 protein, sequestosome-1 or profilin-1 were reported to bind multiple allele-specific MHC class II molecules. In contrast, peptides deriving from microtubule-associated protein tau, presenilin 2 and serine/threonine-protein kinase TBK1 were exclusively reported to bind MHC molecules encoded by the HLA-DRB1 1501 allele, a recently-identified susceptibility gene for late onset Alzheimer’s disease. Finally, we observed that the whole list of proteins reported to provide endogenous MHC class II-binding peptides in human B-cells is specifically enriched in neurodegeneration-associated proteins. Overall, our work indicates that immunization against neurodegeneration-associated proteins might be a physiological process which is shaped, at least in part, by B-cells.

## INTRODUCTION

Multiple studies have now established that neurodegenerative disorders are not cell-autonomous. The pathophysiological processes leading to neurodegeneration involve and target not only neurons but also glial cells, including astrocytes, microglia and oligodendrocytes (1–3). Moreover, beyond central nervous system (CNS) cells, the adaptive immune system has emerged as a potentially important player. When considering only T-cell responses, T-cell reactivity against amyloid beta peptides was already reported more than a decade ago in both patients suffering from Alzheimer’s disease (AD) and aged healthy subjects (4). Recent works further demonstrated that alpha-synuclein (SNCA)-derived peptides elicit helper and cytotoxic T-cell responses in a subgroup of Parkinson’s disease (PD) patients harboring specific major histocompatibility complex (MHC) alleles (5, 6). Similarly, in animal models of neurodegenerative disorders, T-cell responses against peptides deriving from neurodegeneration-associated proteins were also demonstrated. In particular, in the MPTP (1-methyl-4-phenyl-1,2,3,6-tetrahydropyridine) model of PD, pathogenic brain-infiltrating T-cells directed against SNCA were found to target the substantia nigra of diseased mice (7, 8). Along this line, in a mouse model of tauopathy, CD8 cytotoxic T-cells directed against tau protein were shown to infiltrate the hippocampus and to drive cognitive alterations (9). Finally, besides AD and PD, several studies provided evidence that T-cells are activated during the course of Huntington’s disease (HD), amyotrophic lateral sclerosis (ALS) and/or their animal models (10–13). Nevertheless, the putative autoantigens targeted under those conditions remain unknown.

Extending the functional results mentioned above, genome-wide association studies (GWAS) have unraveled the central position of autoimmunity-related genes in the genetic susceptibility landscape of neurodegenerative diseases (14–19). This is notably the case for *HLA-DRB1* alleles which were recently demonstrated to confer increased risks of developing PD (15) or late-onset AD (14). Also, two genes involved in familial forms of PD, namely *PINK1* (PTEN-induced putative kinase 1) and *PRKN* (Parkin RBR E3 Ubiquitin Protein Ligase also named PARK2 or Parkin) were shown to regulate the presentation of mitochondria-derived antigens in the context of MHC class I molecules (20). Finally, innate immune functions were demonstrated for several genes causatively-linked to familial forms of PD or ALS. These comprise *C9ORF72* (21–23), *LRRK2* (Leucine-rich repeat kinase 2) (24–27), *GRN* (Granulin Precursor) (28, 29) as well as *PRKN* and *PINK1* (30–34).

However, advocating for the role of autoimmunity in neurodegenerative disorders requires yet addressing several important issues. In particular, it appears crucial to determine how T-cells directed against neurodegeneration-associated antigens are primed in the periphery. The extent to which autoimmunity against neurodegeneration-associated antigens might be a physiological event needs also to be assessed. Last but not least, a global view on the expression of neurodegeneration-associated proteins by professional antigen-presenting cells (APCs) is still lacking. In an attempt to address these issues, we used here a systems biology approach embracing a large range of previously-published experimental data. We notably explored in normal human APCs the expression patterns and reported protein-protein interactions of the most common neurodegeneration-associated proteins. Overall, our results indicate that in human B-cells, a large majority of neurodegeneration-associated proteins are expressed at the protein level and form a complex network centered on the inflammation/autophagy-related molecules SQSTM1 (Sequestosome-1) and TRAF6 (TNF Receptor Associated Factor 6). Finally, the analysis of MHC class II immunopeptidome databases provides evidence that neurodegeneration-associated proteins expressed by human B-cells are a source of endogenous peptides which are presented in the context of HLA class II molecules.

## MATERIALS AND METHODS

### Workflow

A scheme summarizing the workflow followed in the present work is shown in Figure 1.

**Figure 1:**
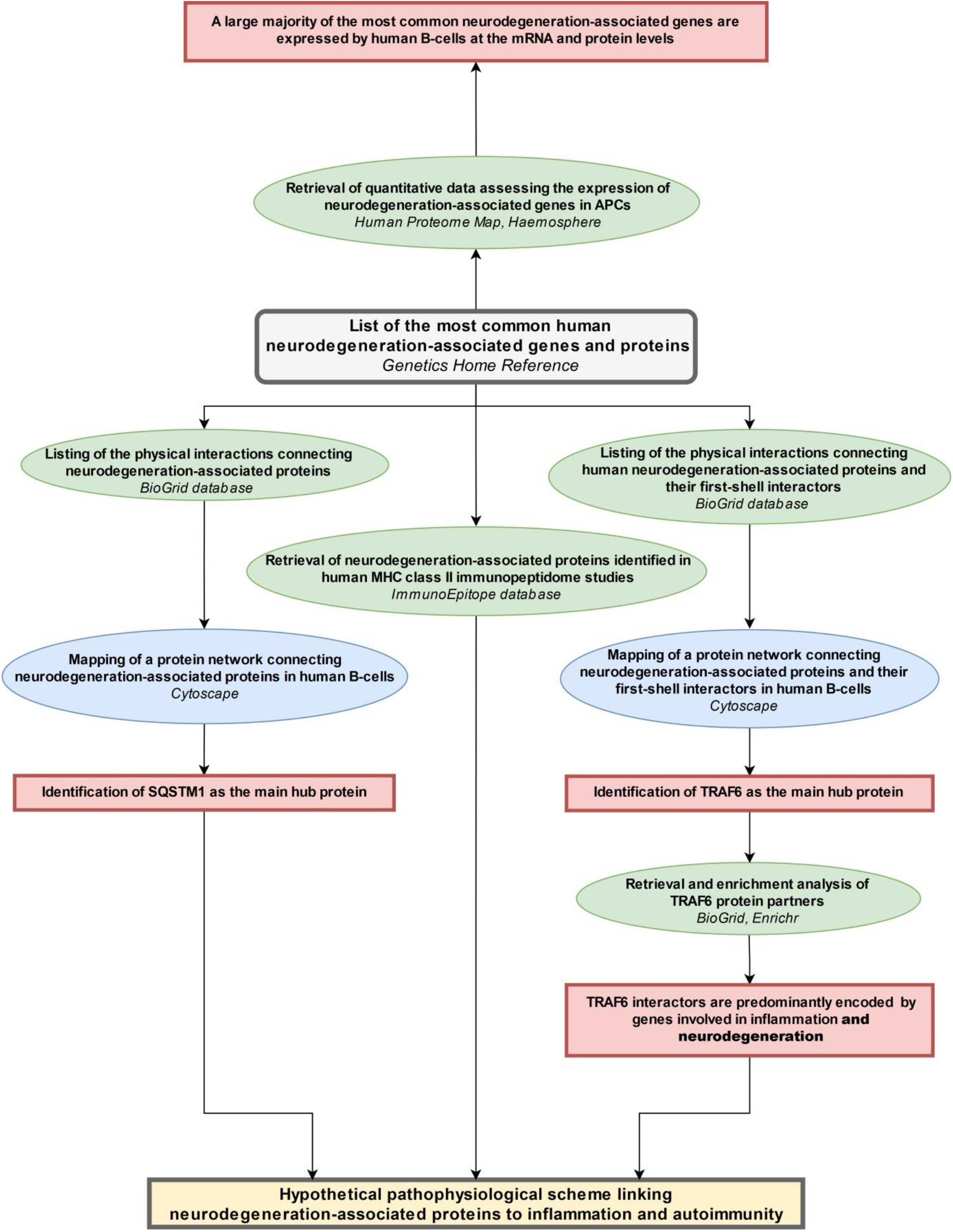
Workflow of the sudy. The workflow starts from the upper central grey rectangle. Other rectangles (yellow or red) frame the main results obtained following each of the analytical steps which are briefly described in ellipse shapes (green or blue). Terms in italics correspond to the name of the bioinformatics tools used for each analytical step. MHC: major histocompatibility complex; APCs: antigen-presenting cells.

### Data mining methods and bioinformatics tools

All the bioinformatics analyses were performed at least 3 times between March 2018 and September 2019. Databases, bioinformatic tools and corresponding tasks performed in this study are described below.

<u>- The Genetics Home Reference website</u> (35), is a regularly updated consumer health resource from the National Library of Medicine. It provides information to the general public about the effects of genetic variation on human health. In the present paper, we used the “Genetics Home Reference” website to select without *a priori* the most common genetic variations/alterations that are linked to the following neurodegenerative disorders: Alzheimer’s disease, Parkinson’s disease, amyotrophic lateral sclerosis, Huntington’s disease, fronto-temporal dementia (FTD).

<u>- The enrichment web platform Enrichr</u> (36) perform enrichment analyses from queried lists of genes. The Enrichr website allows surveying simultaneously 132 libraries gathering 245,575 terms and their associated lists of genes or proteins. Enrichment analysis tools provided by the Enrichr bioinformatics platform provides adjusted P-values computed from the Fisher’s exact test. We focused our analysis on the “Jensen DISEASES” ontology library (37) which is based exclusively on text-mining and allow determining whether a list of genes is significantly associated with specific disease-related terms.

<u>- The BioGrid</u> database (38) compiles 29 169 experimentally-proven protein-protein interactions connecting 23 098 human proteins. Querying the BioGrid database allows retrieving for any given human protein the current list (update in 2019) of published experimentally-identified human protein partners.

<u>- The Human Proteome Map database</u> (39) compiles protein expression data obtained by mass spectrometry from human normal tissues and cells including a total of 85 human blood samples from which the protein profiles of 6 blood-circulating cell types were established. In parallel, we also explored 3 recently-published mass spectrometry datasets reporting on the protemics profiles of blood-derived human B-cells (40), blood-derived human dendritic cells (DCs) (41) and cultured monocyte-derived human dendritic cells (MoDCs)(42).

<u>- The Immune Epitope Database and Analysis Resource (IEDB-AR)</u> (43) compiles experimental data on antibody and T-cell epitopes in humans, non-human primates and other animal species. We followed a 3 step strategy as described below.

#### Step 1: retrieval of a list of human MHC class II binding peptides and their parent proteins

- in the “Epitope” tab, the “Any Epitopes” item was marked
- in the “Assay” tab, the “MHC Ligand Assays” item was marked
- in the “Antigen” tab, “Homo sapiens” was entered in the “Organism” line
- in the “MHC restriction” tab, the “MHC class II” item was marked
- in the “Host” tab, the “Humans” item was marked
- in the “Disease” tab, the “Any Disease” item was marked.

Search was then launched and, from the results page, the “Assays” tab was selected and a list of currently known ligands (peptides and parent proteins) of human MHC molecules was retrieved.

#### Step 2: filtering the results

We retained only results which were both: i) obtained by mass spectrometry analysis of peptides eluted from MHC class II molecules and ii) generated from cells of the B-cell lineage, in the absence of immunization or stimulation protocols.

#### Step 3: checking the reference IDs associated to each identified parent protein

For each selected study and/or set of results, the reference ID associated to each parent protein was checked on the UniProt website (44) in order to avoid redundant or obsolete IDs.

<u>- The Cytoscape software</u> (45) is an open source software allowing to visualize complex networks. We used the function “Network analysis” to identify hubs i.e objects exhibiting the highest number of degrees (connections to other objects) in the generated networks.

### Statistics

When not embedded in bioinformatics webtools, statistics for enrichment analyses were performed using the Fisher exact test. In particular, we assessed for several identified hub proteins whether their retrieved list of partners (obtained via the BioGrid database) was statistically enriched in neurodegeneration-associated proteins. To calculate enrichment factors, we set the expected reference ratio as 0.002 which corresponds to the number of common neurodegeneration-associated proteins (i.e 48 according to the “Home Genetics Reference” website) over the number of coding genes for which interactors can be retrieved from the BioGrid database (i.e 23098). The obtained p-values were then adjusted using Bonferroni correction. The same approach was used to determine whether lists of parent proteins from which derive human MHC class II-binding peptides are significantly enriched in neurodegeneration-associated proteins.

## RESULTS

### 1- Neurodegeneration-associated proteins form a unique interaction network in which PRKN is the main hub

From the most recently updated version of the human proteome compiled in the “BioGrid” database, we extracted the currently published and experimentally-proven protein-protein interactions connecting the most common neurodegeneration-associated proteins. Self-interactions were excluded from our analysis. From these retrieved interactions we were able to build and visualize a protein network which encompasses 35 (72%) of the 48 most common neurodegeneration-associated proteins (Figure 2). Interestingly, network analysis showed that PRKN harbored the highest number of interacting partners (n=17). Moreover, when considering the whole list of known PRKN protein partners, the calculated enrichment factor in neurodegeneration associated protein reached 18.97 and was highly significant (Fisher exact test p-value: 3.35E^-15^). This result points to a yet unnoticed property of PRKN as a major hub protein connecting a large array of proteins involved in not only PD but also ALS, AD, Huntington’s disease and/or FTD.

**Figure 2:**
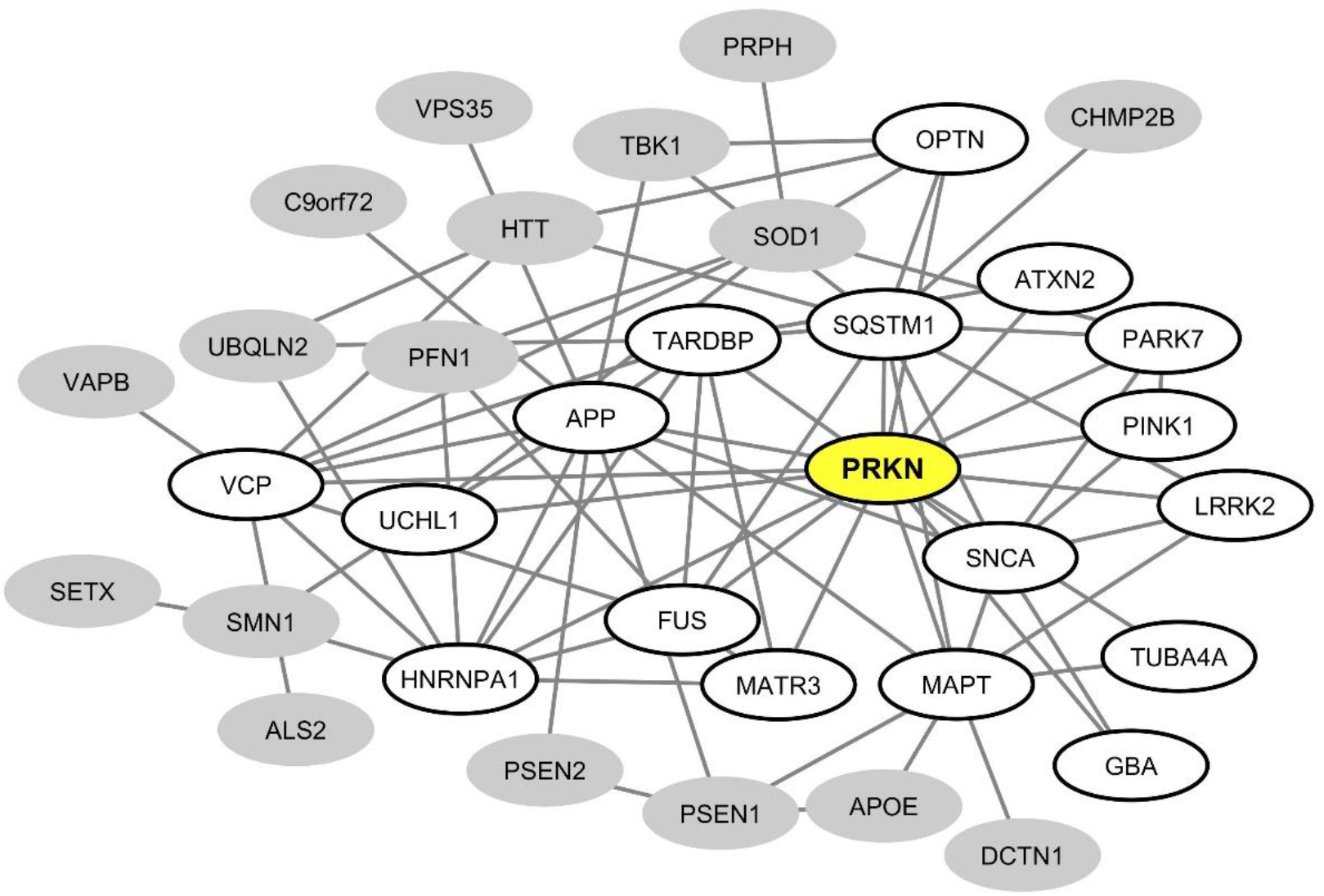
Mapping of the protein-protein interaction network linking common neurodegeneration-associated proteins irrespective of the cell type considered. A survey of the human proteome was performed by querying the protein-protein interaction database BioGRID (38). Each node represents a protein indicated by the corresponding gene symbol and each edge represents an experimentally-demonstrated protein-protein interaction. In this network, the “PRKN” (Parkin) node, highlighted in yellow, exhibits the highest degree (i.e the highest number of direct interactors). White nodes correspond to the first-shell partners of PRKN. Grey nodes are not direct interactors of PRKN.

### 2- A majority of common neurodegeneration-associated proteins are abundantly expressed in human B-cells

We then explored proteomics databases to determine whether neurodegeneration-associated proteins are expressed by APCs under physiological conditions. Our search was restricted to 2 cell lineages which harbor demonstrated antigen-presenting functions: B-cells (46–48) and dendritic cells (DCs) (49–51). Results were combined with those retrieved from proteomics data obtained in 3 independent studies by the mass spectrometry analysis of human B-cells (40) or human dendritic cells (41, 42). Since mass spectrometry is a not a sensitive technique, one may consider that detected proteins are abundantly expressed. As shown in table 1, it is striking that 34 (70%) of 48 common neurodegeneration-associated proteins are detectable by mass spectrometry in human B-cells (Table 1). In human DCs, 20 (41%) of 48 common neurodegeneration-associated proteins were detected and all of these were also detected in human B-cells. Conversely, 14 common degeneration-associated proteins were reported to be detected in B-cells but not DCs. These include notably apolipoprotein E, amyloid beta A4 protein, presenilin-1 and alpha-synuclein.

**Table 1:**
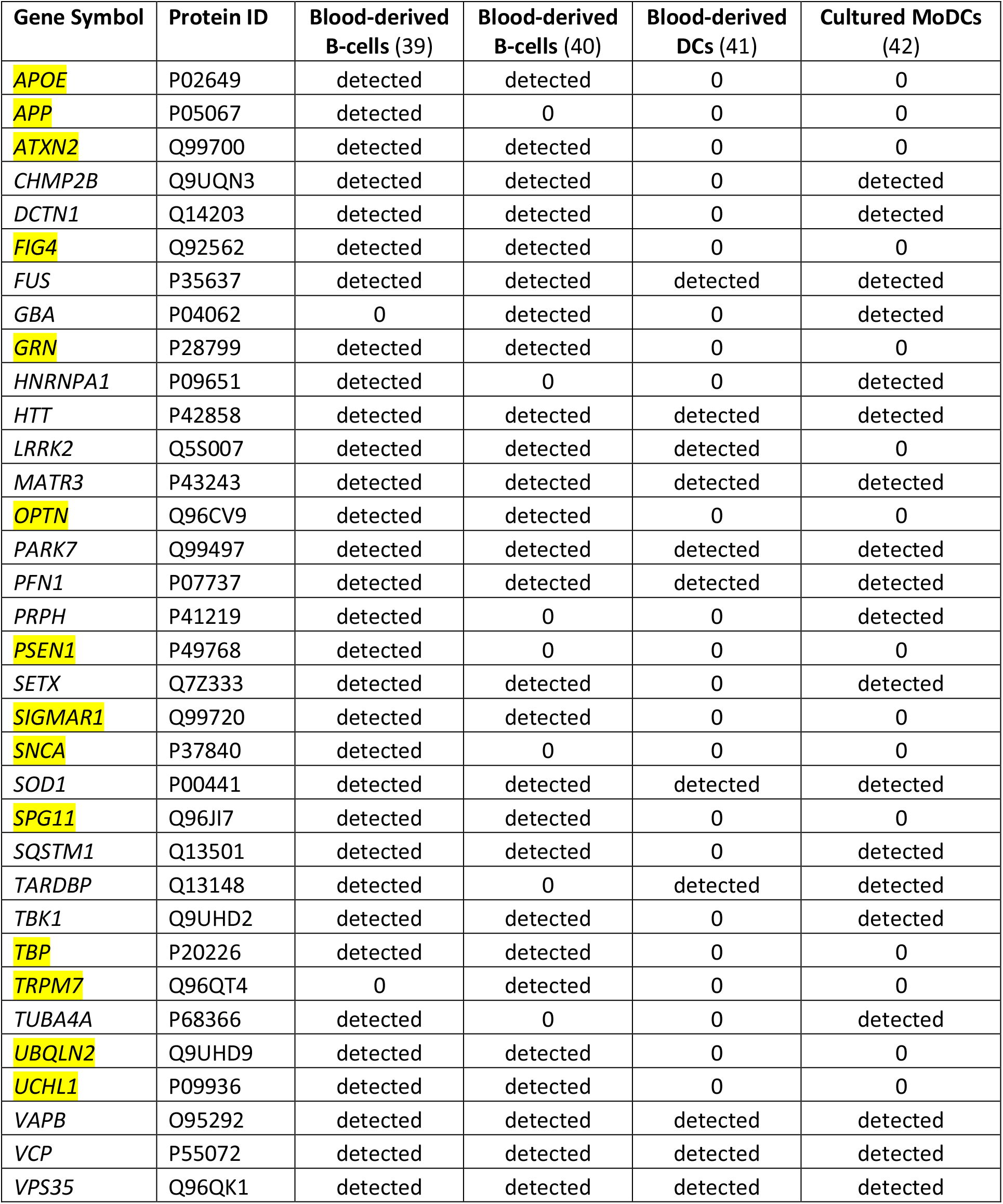
Expression pattern of common neurodegeneration-associated proteins in human B-cells and human dendritic cells. The proteomics profiles of human circulating B-cells, circulating dendritic cells (DCs) or cultured monocyte-derived dendritic cells (MoDCs) were retrieved from 4 independent mass spectrometry studies (39–42). We found that 34 from 48 common neurodegeneration-associated proteins were detected by mass spectrometry in at least 1 of the explored studies. Interestingly, 14 common degeneration-associated proteins (highlighted in yellow) were reported to be detected in B-cells but not DCs. These include notably apolipoprotein E, amyloid beta A4 protein, presenilin-1 and alpha-synuclein. In contrast, when detected in DCs, common neurodegeneration-associated proteins were constantly detected in B-cells.

Of note, in both human B-cells and DCs, *PRKN* was not detected at the protein level by mass spectrometry. This finding urged us to identify the main hub protein(s) which may interconnect abundantly-expressed neurodegeneration-associated proteins in human B-cells.

### 3- Neurodegeneration-associated proteins expressed by human B-cells form a unique interaction network in which the inflammation/autophagy-related protein SQSTM1 is the main hub

From the interaction network depicted in Figure 2 we only retained nodes corresponding to neurodegeneration-associated genes that are detectable by mass spectrometry in human B-cells. We observed that in this cell type, 24 neurodegeneration-associated proteins are predicted to form a tight interaction network which is centered on the inflammation/autophagy-related protein SQSTM1.

### 4- The protein network formed by neurodegeneration-associated proteins connects with a set of first shell interactors among which the B-cell inflammation/autophagy-related protein TRAF6 is the main hub

To extend these data, we then retrieved from the “BioGrid” database the currently known and experimentally-demonstrated interactors of common neurodegeneration-associated proteins, irrespective of their levels of expression in human B-cells. From this list, we extracted candidate hub proteins interacting with more than 10 neurodegeneration-associated proteins. For each of these candidate hubs, total lists of protein interactors were then retrieved from BioGrid and a Fisher exact test was applied so to determine if neurodegeneration-associated proteins were actually significantly enriched in each established list. By this mean, we were able to assign an enrichment factor and an associated p-value to each candidate hub. This approach allowed us to identify 20 hub proteins (Table 2) with which common neurodegeneration-associated proteins are specifically connected. Among these 20 hub proteins, 10 are expressed at the protein level as assessed via the “Human Proteome Map” database (39). Of note, from the 10 hub proteins exhibiting the most significant and highest enrichment factors, 8 are abundantly expressed by human B-cells. These include: i) the heat shock proteins HSPA8 (also named HSC70), HSPA4 (also named HSP70) and HSP90AA1 (also named HSP90) which are all involved in both antigen presentation by MHC class II molecules (52–54) and the modulation of T-cell responses (55–57) and ii) YWHAZ and YWHAQ proteins also known as 14-3-3 protein zeta and theta respectively, which bind MHC class II molecules (58, 59) and are implicated in various neurodegenerative diseases (60–62). Finally, unexpectedly, the most significant hub we identified was TRAF6 (TNF receptor associated factor 6), an inflammation/autoimmunity-related molecule (63, 64) playing a major role in the control of B-cell activation (65, 66). Indeed, the whole list of TRAF6 interactors comprises 17 neurodegeneration-associated proteins which corresponds to an enrichment factor of 27.68 and an adjusted p-value of 1.57E^-16^.

**Table 2.**
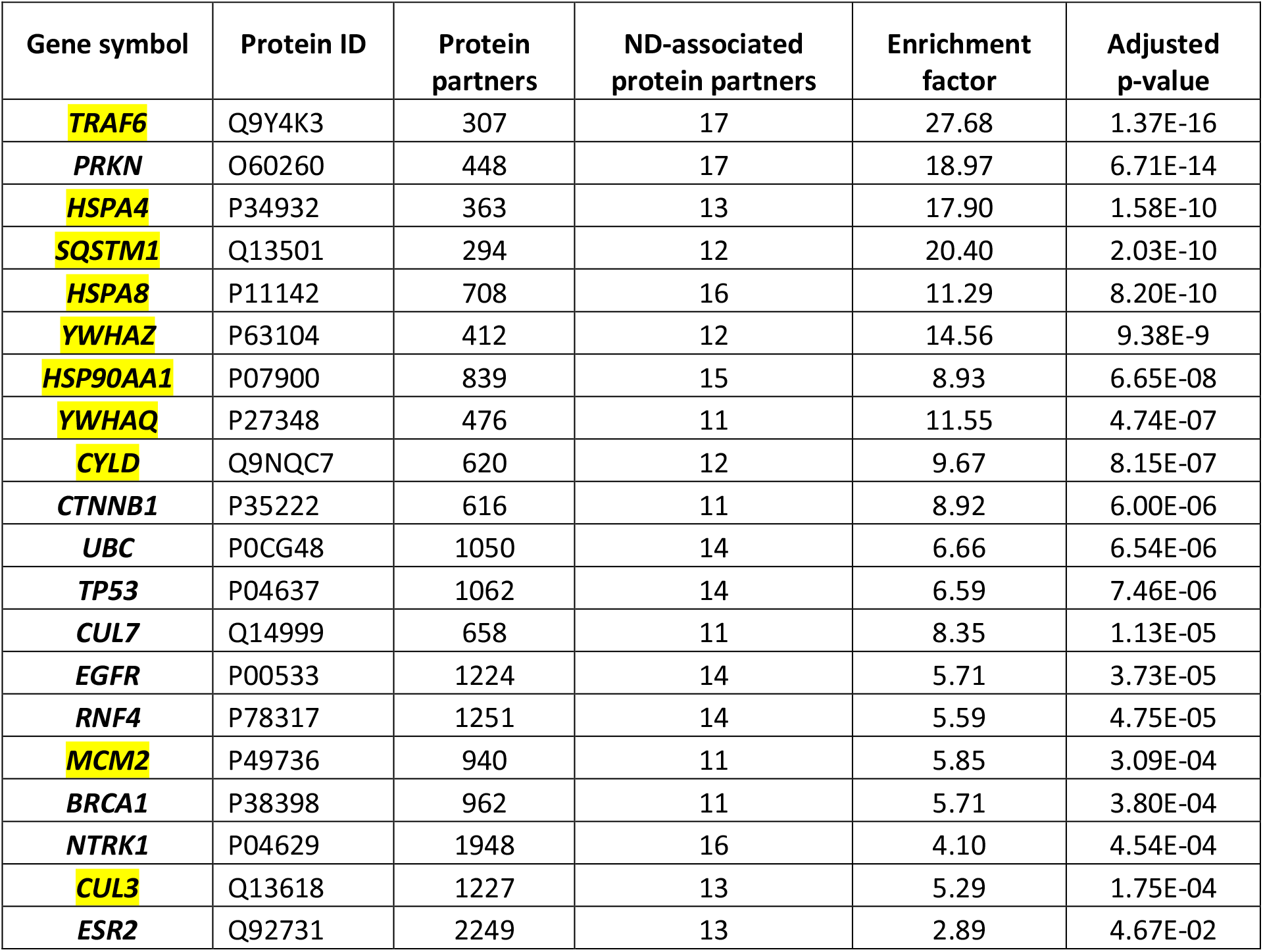
A survey of the human proteome was performed by querying the protein-protein interaction database BioGRID (38). Candidate hub proteins interacting with more than 10 common neurodegeneration-associated proteins were retrieved. The lists of partners currently demonstrated for each of these candidate hubs were then retrieved. For each of these lists, a factor of enrichment in neurodegeneration-associated proteins was established along with an associated Fisher’s exact test p-value, as described in the Materials and Methods section. From left to right, gene symbols and corresponding protein IDs of the identified hub proteins are shown in the first and second column, the associated total numbers of partners and total numbers of neurodegeneration (ND)-associated protein interactors are shown in the third and fourth column. The corresponding enrichment factors and adjusted p-values are shown in the last two columns. Proteins that are detectable by mass spectrometry in human B-cells, according to the “Human Proteome Map” database (39), are highlighted in yellow.

However, one may argue that the list of currently-known TRAF6 partners might be enriched in not only neurodegeneration-associated proteins but also many other sets of proteins which do not relate with neurodegeneration. To address this issue, we performed on the whole list of TRAF6 partners a non *a priori* enrichment analysis using the “JENSEN Disease” text-mining webtool (37) embedded in the “Enrichr” analysis platform (36). Results shown in Table 3 indicate that the 10 disease-related terms with which TRAF6 partners are the most significantly associated comprise the terms “Frontotemporal dementia”, “Neurodegenerative disease” and “Pick’s disease”. This finding points to a specific link between TRAF6 and neurodegeneration.

**Table 3.**
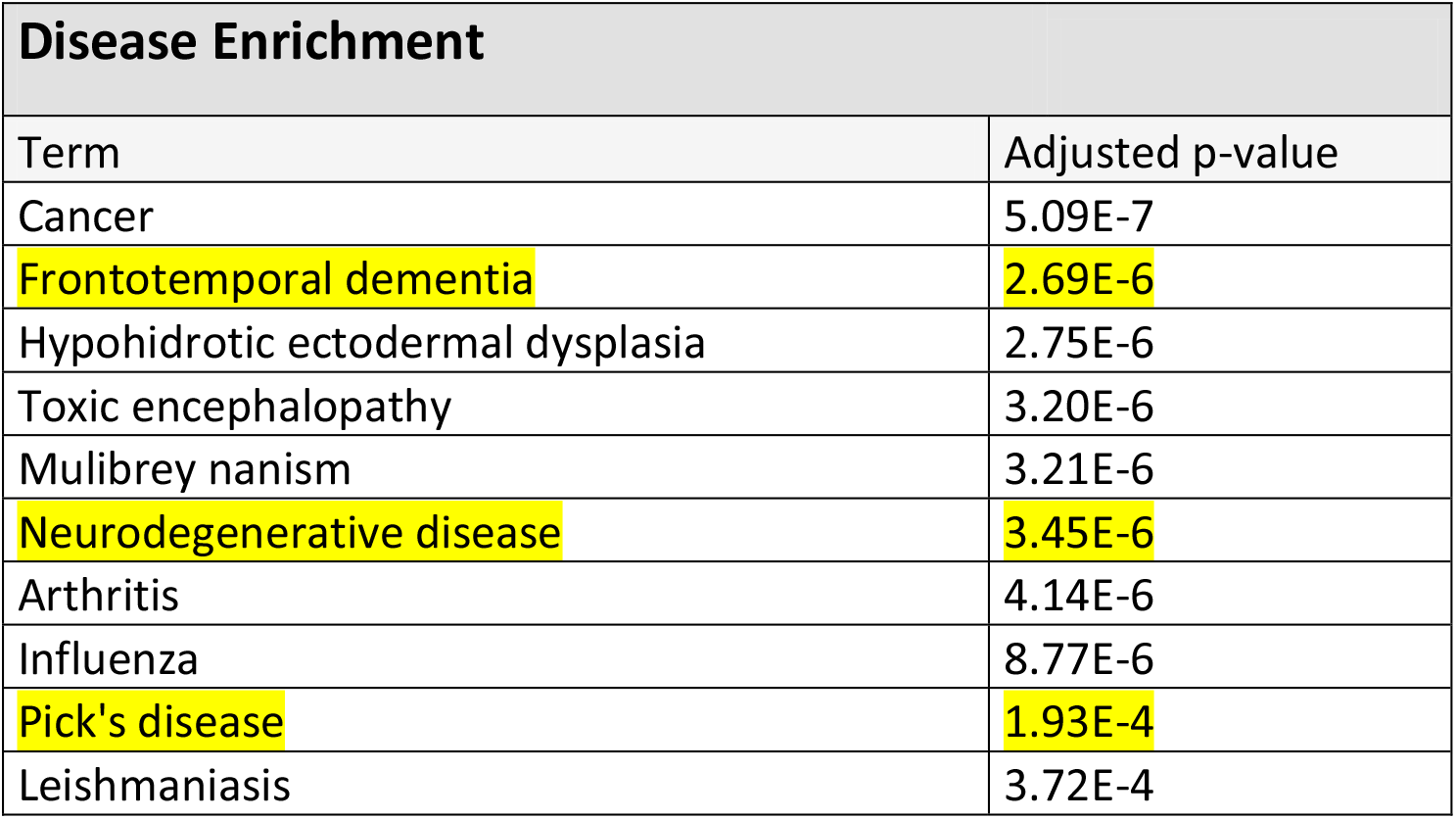
A survey of the human proteome was performed by querying the protein-protein interaction database BioGRID (38). The list of currently known and experimentally-demonstrated TRAF6 partners was retrieved. From this list of interactors, an enrichment analysis with the “JENSEN Disease” text-mining webtool (37) embedded in the “Enrichr” analysis platform (36) was then performed. Shown are the disease-related terms exhibiting the 10 most statistically significant enrichments. Terms relating with neurodegeneration are highlighted in yellow.

As a control and to further establish the specificity of our findings, we assessed whether similar results would be obtained from a list of common demyelination-associated proteins i.e. proteins commonly considered as candidate autoantigens in multiple sclerosis (67). We found that neither TRAF6 nor the other hubs linking common neurodegeneration-associated proteins exhibited lists of protein partners which were significantly enriched in common demyelination-associated proteins (data supplement 1).

From these results we then built and visualized a B-cell protein-protein interaction network encompassing the most significant hub proteins and their neurodegeneration-associated interactors expressed in B-cells (Figure 5). We observed that from the 32 neurodegeneration-associated proteins expressed by B-cells, 22 (68%) are first shell interactors of HSPA4, HSPA8, TRAF6 or SQSTM1. These results unravel a yet unknown function of these molecules as major hub proteins connecting in B-cells a large array of proteins involved in PD, ALS, AD, Huntington’s disease and/or FTD.

**Figure 5:**
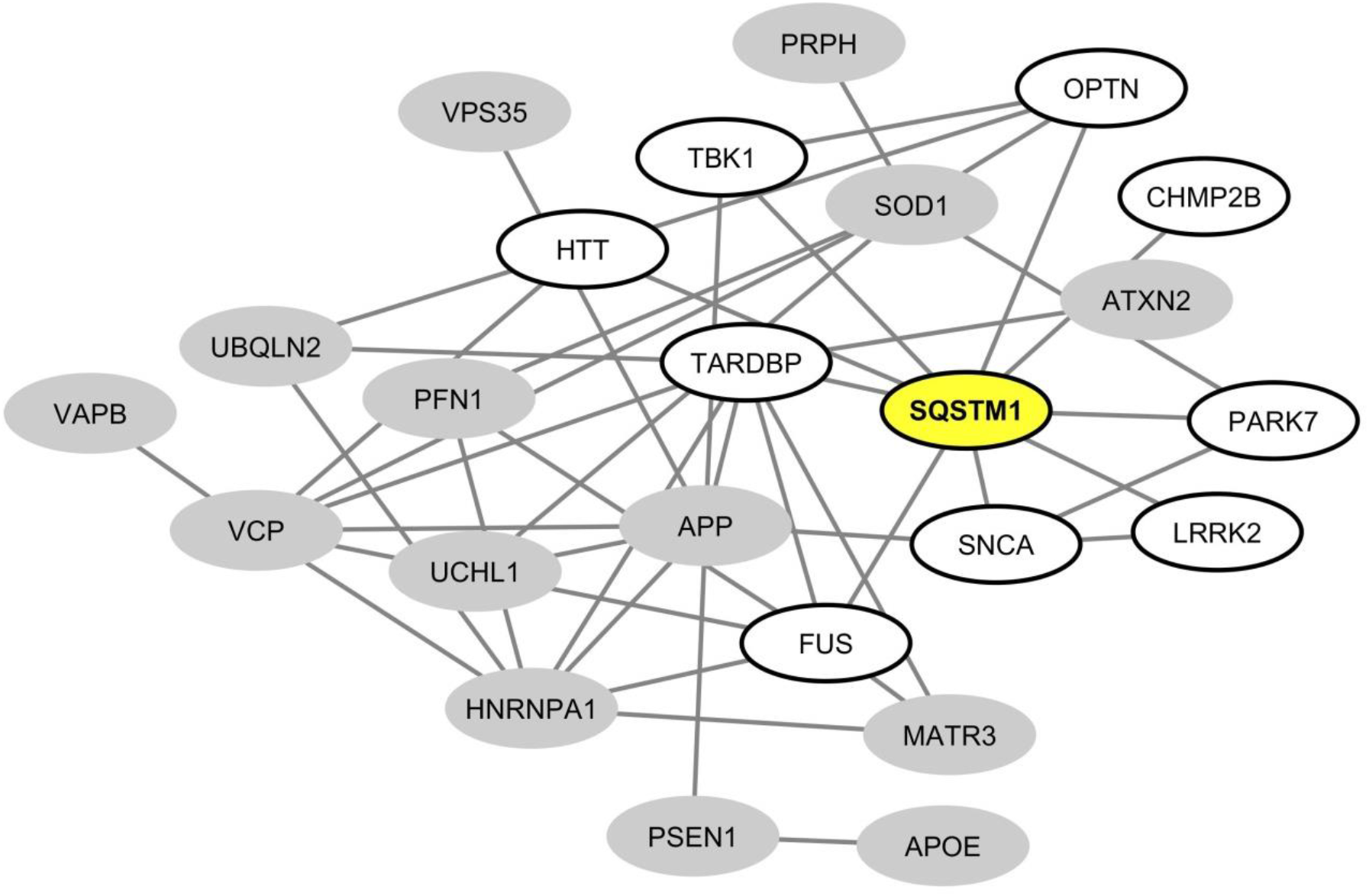
Mapping of the protein-protein interaction network linking common neurodegeneration-associated proteins in human B-cells. A survey of the human proteome was performed by querying the protein-protein interaction database BioGRID (38). Each node represents a protein indicated by the corresponding gene symbol and each edge represents an experimentally-demonstrated protein-protein interaction. In this network, the “SQSTM1” (Sequestosome-1) node, highlighted in yellow, exhibits the highest degree (i.e the highest number of direct interactors). White nodes correspond to the first-shell partners of SQSTM1 in human B-cells. Grey nodes are expressed by B-cells but are not direct interactors of SQSTM1.

**Figure 6:**
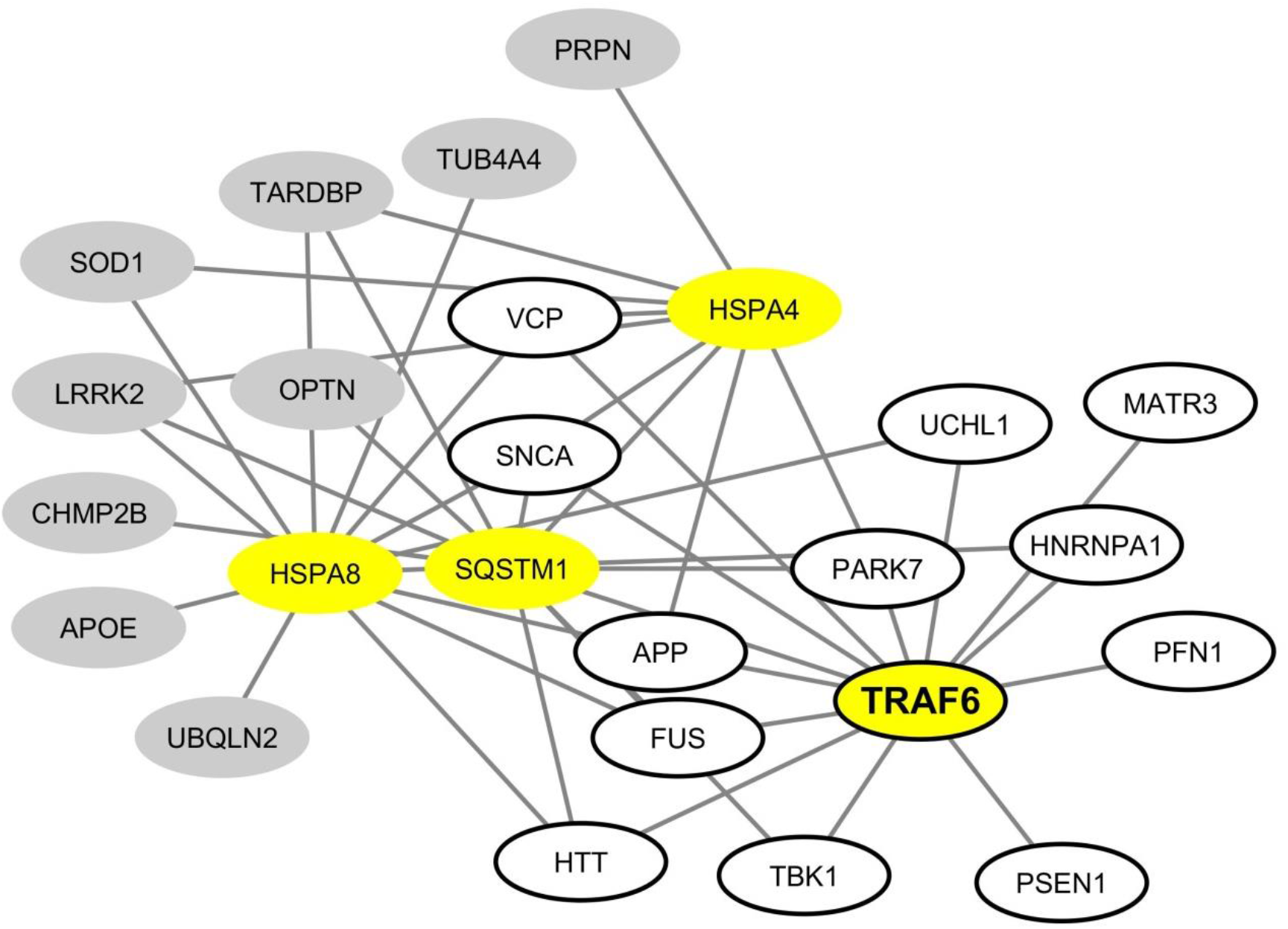
Mapping of the protein-protein interaction network linking common neurodegeneration-associated proteins and their hub protein partners in human B-cells. A survey of the human proteome was performed by querying the protein-protein interaction database BioGRID (38). Only proteins expressed by human B-cells according to the “Human Proteome Map” database (39) were taken into account. Each node represents a protein indicated by its corresponding gene symbol. The 4 hub proteins expressed by B-cells and whose partners exhibited the most significant and highest enrichment factors in neurodegeneration-associated proteins are highlighted in yellow. The neurodegeneration-associated proteins expressed by B-cells and interacting with TRAF6 are shown in white ellipses. Other neurodegeneration-associated proteins involved in this network are shown in grey ellipses.

### 5- Common neurodegeneration-associated proteins provide a source of endogenous MHC class II-binding peptides in human B-cells

While peptides presented by MHC class II molecules are classically generated by the proteolysis of phagocytized exogenous antigens, the presentation of endogenous peptides by MHC class II molecules is an alternate pathway which has been robustly documented (68–71). We thus surveyed the Immune Epitope DataBase (IEDB-AR) (43) to assess whether peptides derived from neurodegeneration-associated proteins had been previously identified as binding MHC class II molecules in immunopeptidome studies which used human cells of the B-cell lineage as a source of endogenous peptides. Of note, we excluded studies assessing the MHC binding of exogenously-provided specific peptides and retained only works relying on the systematic mass spectrometry-based identification of peptides eluted from MHC class II molecules. In addition, we excluded experiments in which immunization or stimulation protocols were applied to B-cells. On this basis, we retained 19 studies (data supplement 2) which were performed on cells of the B-cell lineage including predominantly Epstein-Barr virus (EBV)-transformed B-cells. When screening these studies, we found that 23 out of 48 common neurodegeneration-associated proteins were reported to provide endogenous MHC class II-binding peptides in B-cells (Table 4 and data supplement 2). The most frequently identified neurodegeneration-associated parent proteins are abundantly expressed in B-cells and comprise notably the proteins encoded by: i) the AD-related genes *APP* and *PSEN1*, ii) the ALS/FTD-related genes *PFN1*, *SQSTM1*, *GRN*, *SOD1* and *VCP* and iii) the PD-related proteins *PARK7* and *GBA* (Figure 1).

**Table 4.**
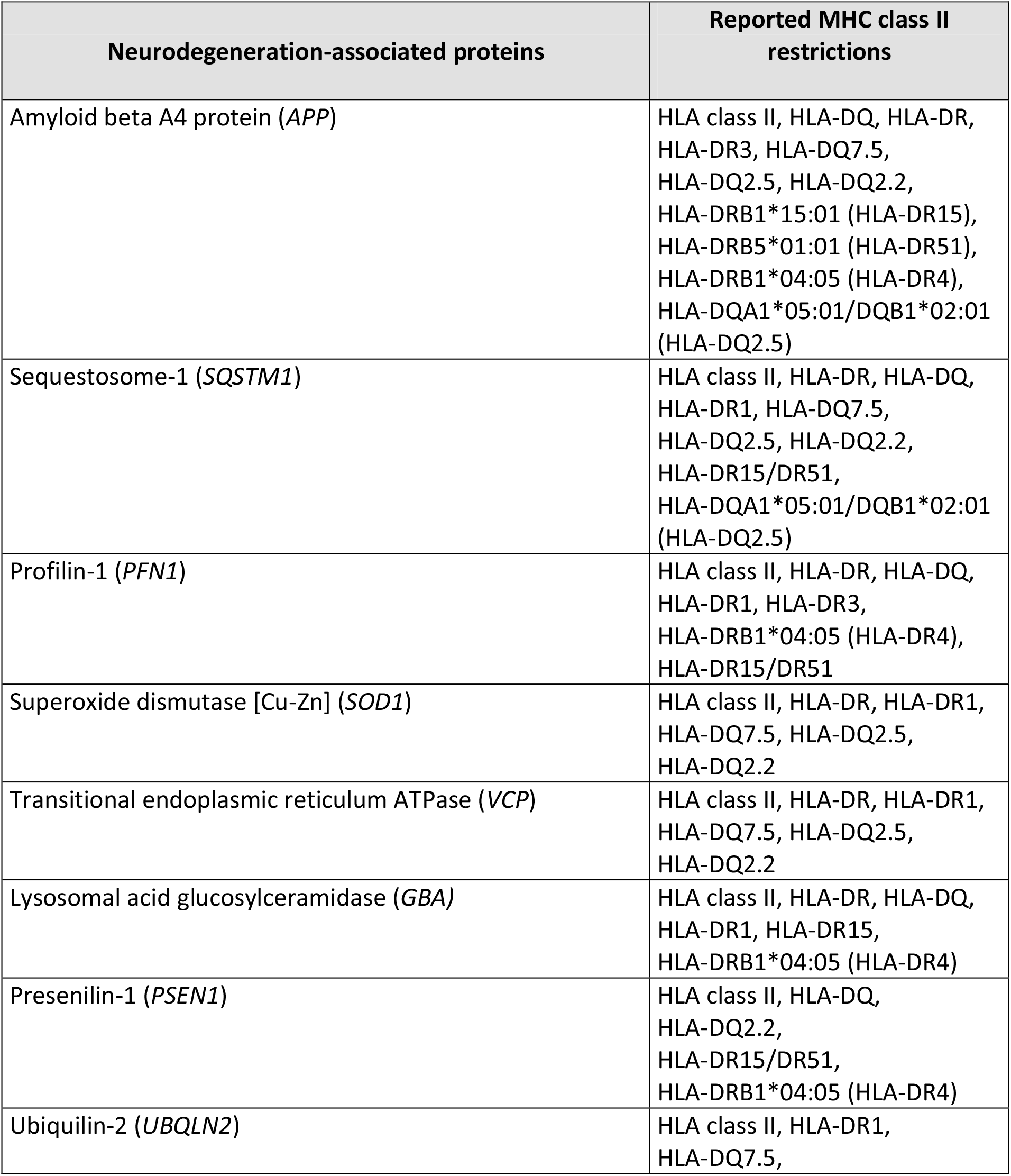
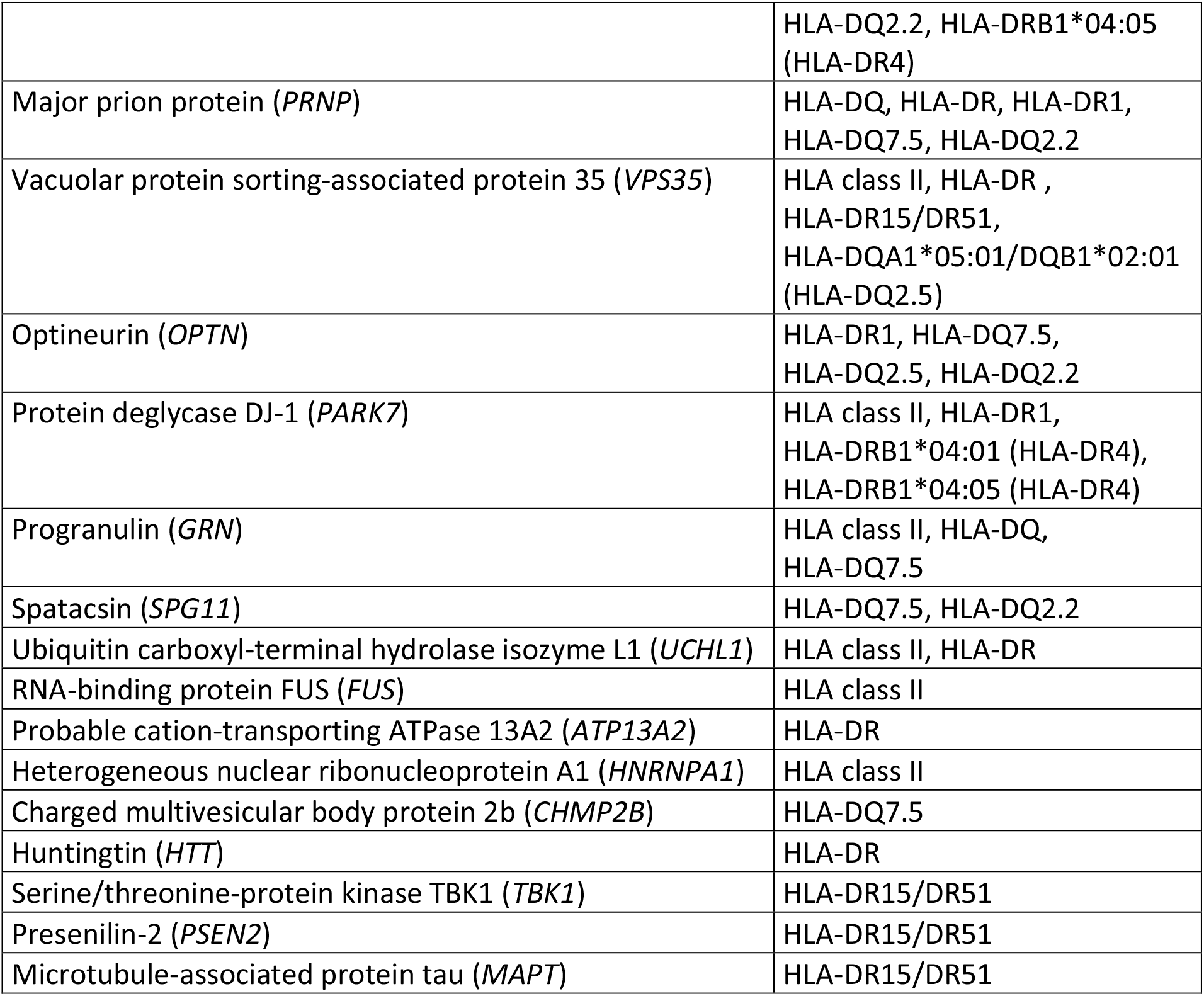
The “IEDB-AR” immune epitope database (43) was screened in order to retain only publications reporting on the systematic mass spectrometry identification of endogenous peptides binding MHC-class II molecules in the human B-cell lineage. Only results obtained on human cells of the B-cell lineage in the absence of immunization or stimulation protocols were taken into account. The retrieved list of parent proteins and corresponding reported MHC class II restriction of derived peptides was then crossed with the list of 48 common neurodegeneration-associated proteins we established. The protein names and gene symbols are shown in the left column; the corresponding reported MHC class II restrictions of derived peptides are shown in the right column. When needed, aliases frequently used in the HLA class II genotype/serotype nomenclature are indicated in brackets.

### 6- Hub molecules linking common neurodegeneration-associated proteins provide a source of endogenous MHC class II-binding peptides in human B-cells

From the 19 relevant B-cell studies we retained on IEDB-AR, we also attempted to determine whether the hub molecules we identified as linking neurodegeneration-associated proteins in B-cells (Table 2) were, in parallel, reported to provide endogenous ligands for MHC class II molecules in B-cells. We found that from 10 candidate hubs abundantly expressed by B-cells, 8 were reported to provide endogenous ligands for MHC class II molecules in human B-cells (Table 5 and data supplement 2).

**Table 5.**
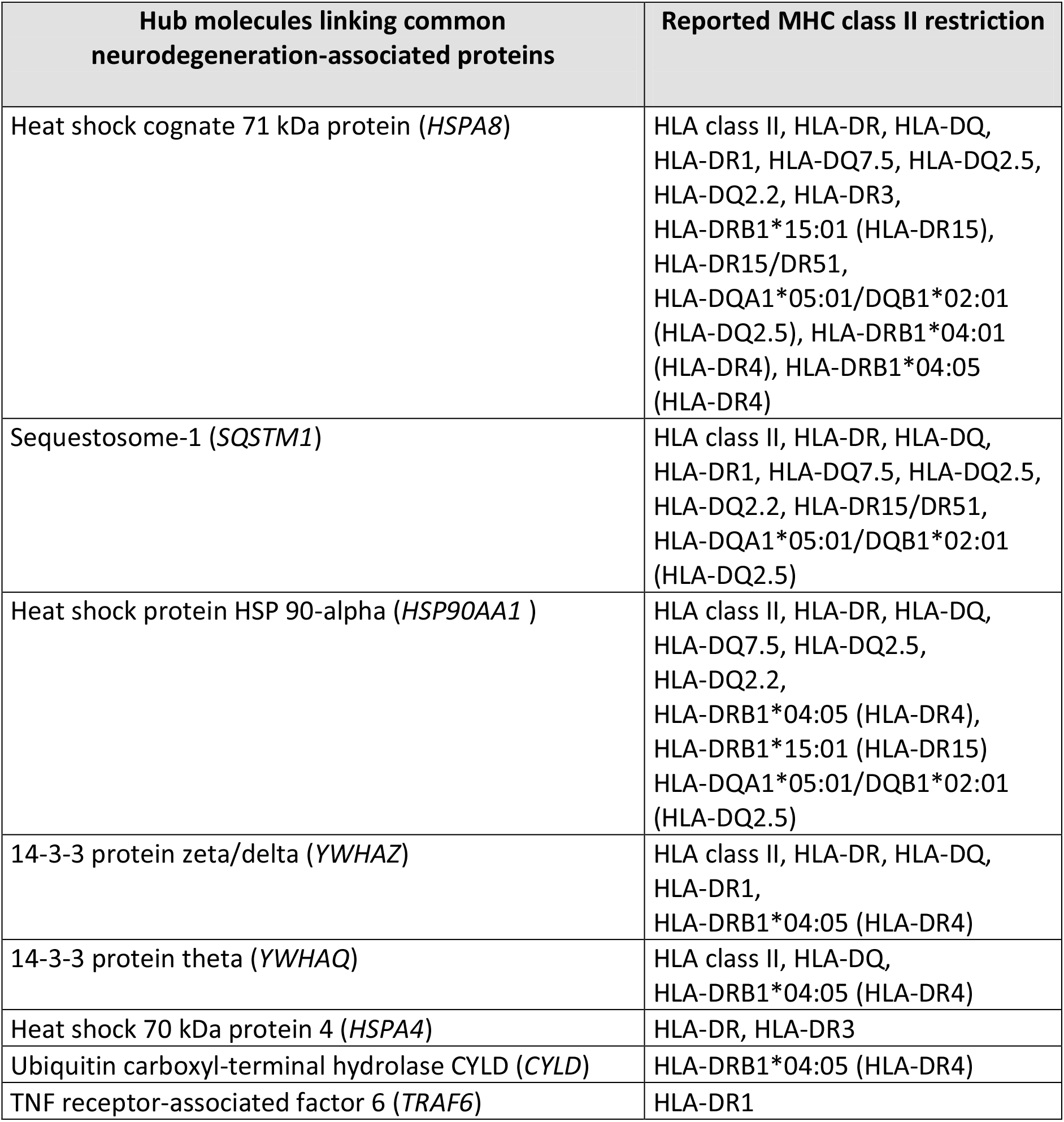
The “IEDB-AR” immune epitope database (43) was screened in order to retain only publications reporting on the systematic mass spectrometry identification of endogenous peptides binding MHC-class II molecules in the human B-cell lineage. Only results obtained in the absence of immunization or stimulation protocols were taken into account. The retrieved list of parent proteins and corresponding reported MHC class II restriction of derived peptides was then crossed with the list of 11 hub molecules being abundantly expressed by human B-cells and linking common neurodegeneration-associated. The protein names and corresponding gene symbols are shown in the left column; the corresponding reported MHC class II restrictions of derived peptides are shown in the right column. When needed, aliases frequently used in the HLA class II genotype/serotype nomenclature are indicated in brackets.

Among these 8 hubs, HSPA8 was the parent protein being the most frequently identified as providing endogenous peptides which bind MHC class II molecules in human B-cells (Table 5).

### 7- In human B-cells, the list of parent proteins from which derive endogenous MHC class II-binding peptides is specifically enriched in neurodegeneration-associated proteins

The whole list of parent proteins identified as providing MHC class II-binding endogenous peptides in B-cells was retrieved from IEDB-AR and an enrichment analysis was performed to determine whether such a list was significantly enriched in neurodegeneration-associated proteins. We found that genes coding for neurodegeneration-associated proteins encompassed 0.62% (22 out of 3523) of the whole genes coding for such parent proteins (data supplement 3), which corresponds to an enrichment of 2.58 (p-value = 0.0006, Fisher exact test) when considering the whole number of human protein-coding genes as roughly 20 000 (72). To confirm these results and assess their level of specificity, we used the “JENSEN Disease” text-mining webtool (37) and performed an unsupervised enrichment analysis from the whole list of genes coding for parent proteins previously identified as providing MHC class II-binding peptides in human B-cells (data supplement 3). From this list of 3522 genes, 96 (2.72%) were annotated with the term “Neurodegenerative disease” (data supplement 3) which corresponds to the second most significant enrichment, after the term “Arthritis” (Table 6).

**Table 6.**
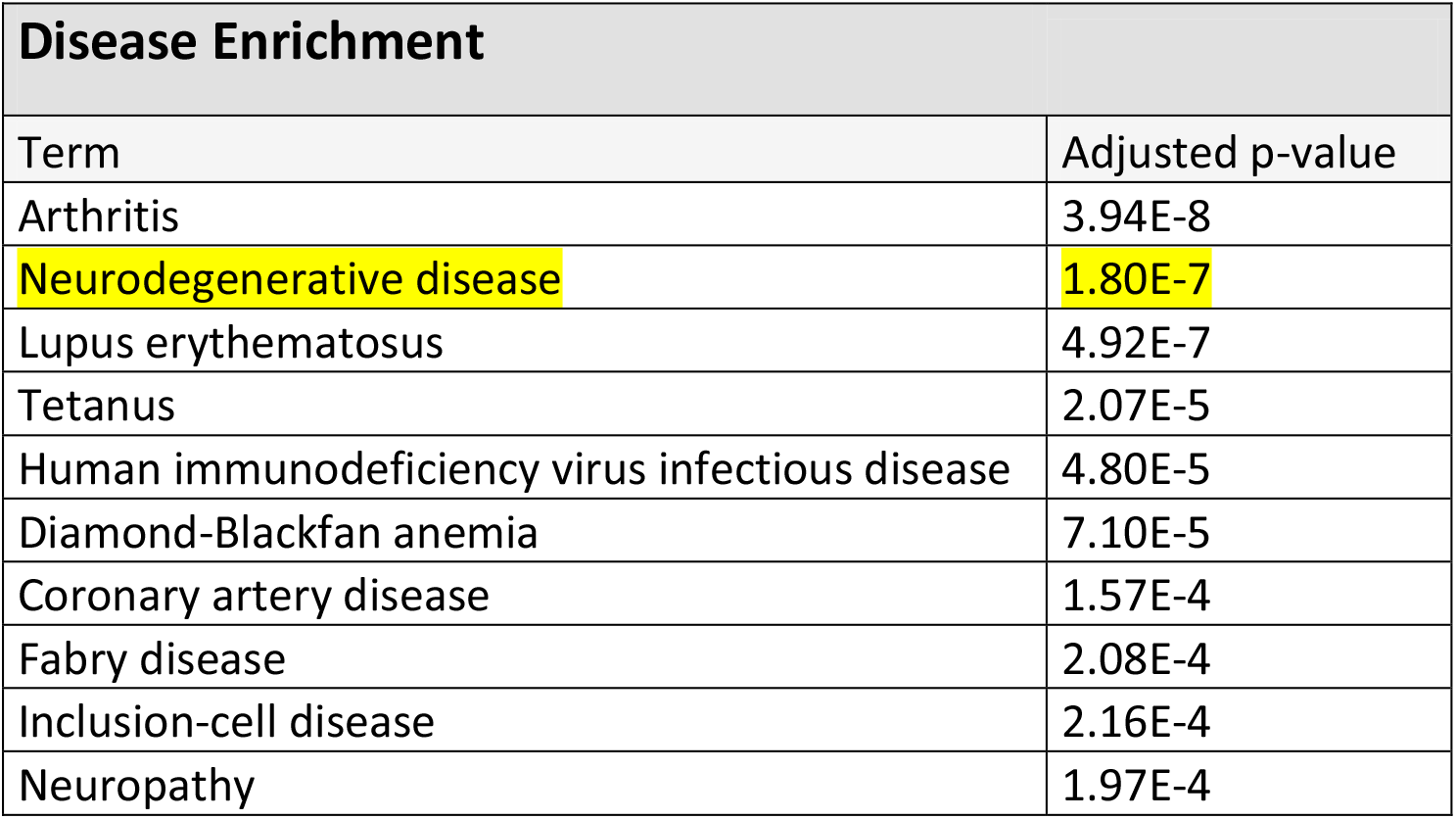
The “IEDB-AR” immune epitope database (43) was screened in order to retain only publications reporting on the systematic mass spectrometry identification of endogenous peptides binding MHC class II molecules in the human B-cell lineage. Only results obtained in the absence of immunization or stimulation protocols were taken into account. The parent proteins from which derive the identified bound peptides were retrieved and the list of corresponding coding genes was submitted to an enrichment analysis with the “JENSEN Disease” text-mining webtool (37) embedded in the “Enrichr” analysis platform (36). The 10 most significant terms associated with this list of genes are shown. The second most significant enrichment is observed for the term “Neurodegenerative disease” (term and corresponding adjusted p-value highlighted in yellow).

Since the HLA-DRB1 1501 allele (corresponding to the HLA-DR15 serotype) was recently identified as a risk factor for sporadic forms of late onset AD (14), we retrieved the whole list of parent proteins (and corresponding coding genes) which, in human B-cells, were previously reported to provide peptides that bind HLA-DRB1 1501-encoded MHC class II molecules. Importantly, such a list was significantly enriched in neurodegeneration-associated proteins (enrichment factor: 5.34; p-value = 0.0003, Fisher exact test). Moreover, when this list of parent proteins was submitted to an unsupervised enrichment analysis, the term “Neurodegenerative disease” was found to reach the highest level of statistical significance (Table 7).

**Table 7.**
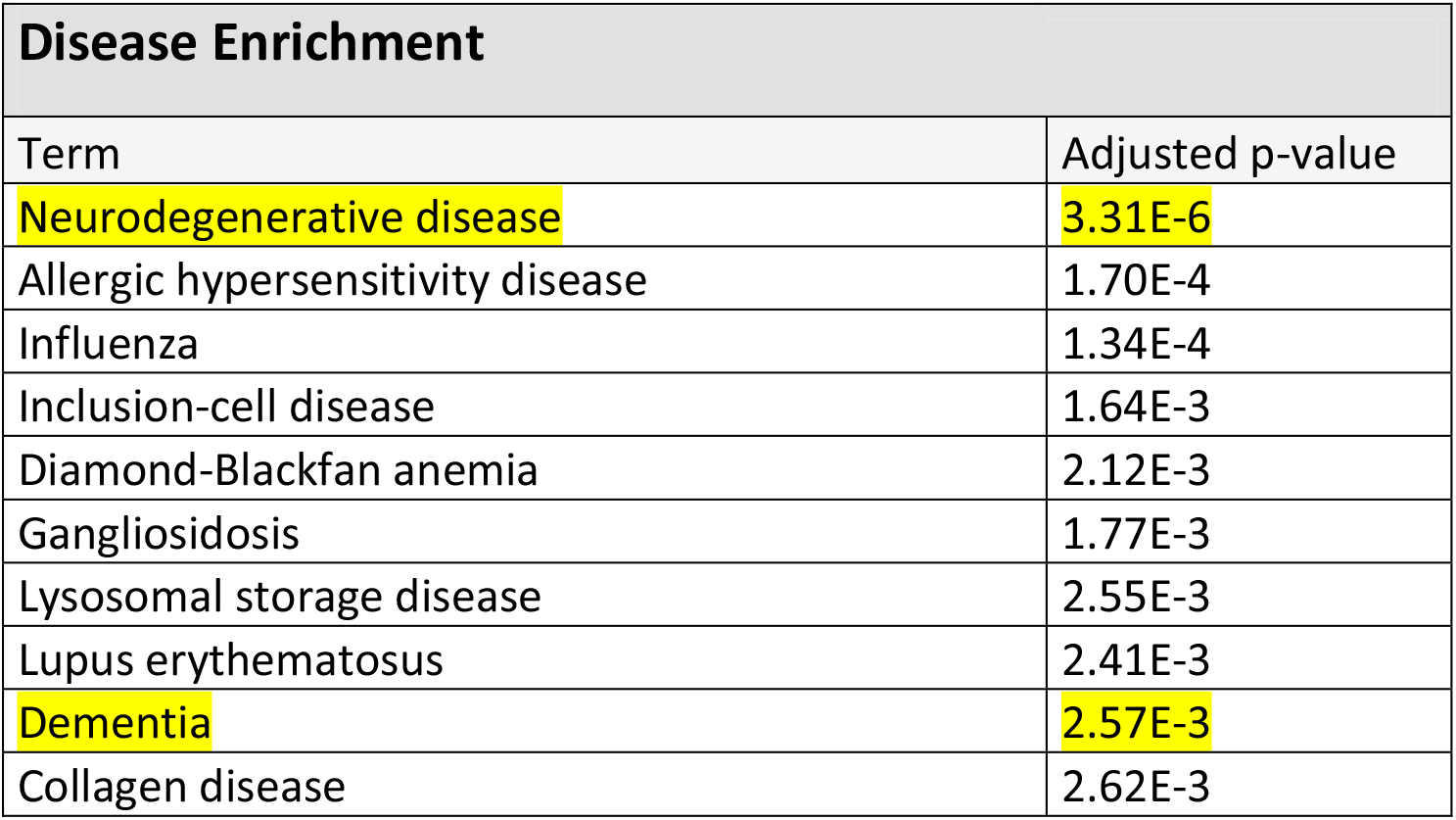
The “IEDB-AR” immune epitope database (43) was screened in order to retain only publications reporting on the systematic mass spectrometry identification of endogenous peptides which bind HLA-DRB1 1501-encoded MHC class II molecules in the human B-cell lineage. Only results obtained in the absence of immunization or stimulation protocols were taken into account. The parent proteins from which derive the identified bound peptides were retrieved and the list of corresponding coding genes was submitted to an enrichment analysis with the “JENSEN Disease” text-mining webtool (37) embedded in the “Enrichr” analysis platform (36). Shown are the disease-related terms exhibiting the 10 most statistically significant enrichments. Terms relating with neurodegeneration are highlighted in yellow.

Finally, based on the IEDB-AR survey we performed, HLA-DRB1 1501-encoded MHC class II molecules are the only HLA class II molecules which, in human B-cells, were reported to bind endogenous peptides deriving from microtubule-associated protein tau, presenilin-2 or serine/threonine-protein kinase TBK1 (Table 4 and data supplement 2).

## DISCUSSION

In the present work, we mined large publically-available databases to provide experiment-based evidence of a link between neurodegeneration and autoimmunity. Using a systems biology approach we report that a large range of common neurodegeneration-associated proteins: i) are expressed by human B-cells under physiological conditions, ii) form a comprehensive and functionally-relevant protein-protein interaction network in human B-cells and iii) provide endogenous peptides which bind MHC class-II molecules in human B-cells. Patients suffering from neurodegenerative conditions exhibit T cell- and/or antibody-mediated responses directed against major neurodegeneration-associated proteins such as amyloid beta A4 protein, alpha-synuclein and tau protein (4–6,73). However, naturally-occurring antibodies against amyloid-beta (74), alpha-synuclein (75–77) and tau protein (73,77,78) were also demonstrated in cohorts of healthy subjects. Similarly, apart from any pathological context, robust T-cell responses against peptides deriving from tau protein were recently demonstrated to widely occur in the general population (79).These findings suggest that the autoimmune processes described in patients with neurodegenerative conditions might be shaped by pre-existing physiological autoimmune responses directed against common neurodegeneration-associated proteins. It is worth noting that, while physiological autoimmunity was firmly demonstrated nearly 50 years ago (80–82), the intimate nature of the links bridging physiological autoimmunity to its pathological counterpart is still matter of debate. Numerous functions have been assigned to physiological autoimmunity (83, 84), including, more recently, a support to cognition (85–90). In line with these findings, our data indicate that a specific set of brain antigens expressed by B-cells and involved in neurodegenerative diseases might prime a neuroprotective and, possibly, cognition-promoting T-cell response under physiological conditions. Of note, B-cells are now recognized as professional APCs and, as such, are able to prime naïve CD4 T-cells with a similar efficiency than DCs (91, 92). The autophagy of cytosolic and nuclear proteins in B-cells was shown to provide a continuous source of endogenous MHC class-II ligands (93). Moreover, such autophagosome-derived peptides induce the proliferation of autologous T-cells under *in vitro* conditions (94). However, that neurodegeneration-associated proteins provide MHC class II-binding endogenous peptides in B-cells neither prove that T-cells are actually primed against such peptides *in vivo* nor that peptide deriving from misfolded neurodegeneration-associated proteins are presented by B-cells under neurodegenerative conditions. Moreover, even if it was actually the case, the phenotype of autoreactive T-cells generated via such a mechanism would need to be explored. More generally, one has to keep in mind that no consensus has been currently reached regarding the phenotype of physiological autoreactive T-cells. Thus, both autoreactive Tregs and autoreactive pro-inflammatory T-cells belong to the physiological T-cell repertoire and were both found to exert neuroprotective effects (95–98). Finally, the picture might be even more tangling since Tregs were demonstrated to prevent autoimmunity by restraining autophagy in APCs (99).

Several genes involved in familial forms of neurodegenerative disorders exert key functions in the autophagy pathway. These notably comprise PRKN, PINK1 and SQSTM1 (100–103). In the recent years, major works provided evidence that, in neurons and immune cells, functional defects in such genes hamper mitophagy (a specialized form of autophagy), stimulate the inflammasome pathway and foster the presentation of mitochondrial antigens by MHC class I molecules (33,104,105). However, these findings do not render account for the existence of HLA class II-restricted T-cell responses against neurodegeneration-associated proteins. Moreover, antigens targeted by autoimmunity during neurodegenerative conditions are far from deriving only from the mitochondrial compartment. In this regard, our work suggests that in B-cells, the inflammation/autophagy-related molecules SQSTM1 and TRAF6 are crucially involved in the presentation of neurodegeneration-associated antigens by MHC class II molecules.

The mining of previously published mass spectrometry analyses performed on MHC class II-eluted peptides showed that, in human B-cells, several neurodegeneration-associated proteins provide endogenous peptides which bind a large range of MHC class II alleles. This is notably the case for amyloid beta A4 protein, sequestosome-1 and profilin-1. Similarly, peptides deriving from HSPA8, a hub molecule which links a high number of neurodegeneration-associated proteins, bind multiple MHC class II alleles. Our observations suggest that these 4 molecules are likely to elicit immune responses in a large range of the human population. Determining whether or not such antigens are targeted by cognition-promoting autoimmunity is a potentially important issue. On another hand, in human B-cells, several neurodegeneration-associated antigens appear to provide endogenous peptides harboring an allele-specific MHC class II restriction in B-cells. For e.g. endogenous peptides deriving from microtubule-associated protein tau, PSEN2 and Serine/threonine-protein kinase TBK1 were exclusively reported to bind HLA-DRB1 1501-encoded MHC class II molecules under the experimental conditions described above. Since the HLA-DRB1 1501 allele is a susceptibility gene for late onset AD (14), this may prove to be of interest in the context of AD pathophysiology. However, additional studies are clearly needed in order to establish, under strictly comparable conditions, the endogenous immunopeptidomes of distinct MHC class II alleles in B-cells. Moreover, it should be reminded that the endogenous immunopeptidome of MHC class II molecules does not match the corresponding repertoire of exogenous peptides (106). Since a widespread T-cell reactivity against Tau peptides was recently demonstrated in the general human population (79), one may anticipate that individuals bearing the HLA-DRB1 1501 allele exhibit functional specificities regarding their anti-Tau T-cell responses.

We previously proposed that the genetic polymorphism of the HLA-DRB1 locus, which, among primates, is extremely high in the human species (107), might allow the allele-specific presentation of distinct sets of “brain superautoantigens” (108, 109) leading, in turn, to the development and maintenance of distinct sets of cognition-promoting T-cells. The present work indicates that common neurodegeneration-associated proteins might represent an important share of brain superautoantigens. Interestingly, recent magnetic resonance imaging studies reported that, in healthy subjects, specific HLA-DR alleles correlate with the volume ranges of specific brain structures (110, 111). Future studies should be designed to determine whether HLA-DR polymorphism might match both brain structural features and the diversity of T-cell responses against common neurodegeneration-associated proteins. Finally, in the human B-cell lineage, MHC class II-binding endogenous peptides are highly significantly and specifically enriched in peptides deriving from common neurodegeneration-associated proteins. This result raises the intriguing possibility that a main function of physiological autoimmunity could be to control the blood-circulating levels of aggregated forms of amyloid beta A4 protein, alpha-synuclein, tau protein and possibly other neurodgeneration-associated proteins.

## Acknowledgments

no specific funds were used for this study. SN performed the bioinformatics analyses and wrote the paper. MG and LP performed quality controls of bioinformatics analyses and wrote the paper. We thank Céline Auxenfans and Pascale Pascal from the “Bank of Tissues and Cells” of the “Hospices Civils de Lyon” for leaving us the opportunity to develop a bioinformatics research group. This paper can be freely accessed via the BioRxiv platform at the following URL: http://biorxiv.org/cgi/content/short/690677v1

