## Supplementary material for "Common neurodegeneration-associated proteins are physiologically expressed by antigen-presenting cells and are interconnected via the inflammation/autophagy-related proteins TRAF6 and SQSTM1": data supplement 1

| **Gene Symbol** | **Protein ID** |
| --- | --- |
| *CLDN11* | O75508 |
| *CNP* | P09543 |
| *MAG* | P20916 |
| *MBP* | P02686 |
| *MOBP* | Q13875 |
| *MOG* | Q16653 |
| *PLP1* | P60201 |

### Legend: A list of 7 common demyelination-associated proteins which were previously identified as putative autoantigens in multiple sclerosis patients was established on the basis of an authoritative review paper by Ben-Nun A et al. (“From classic to spontaneous and humanized models of multiple sclerosis: impact on understanding pathogenesis and drug development. J Autoimm 2014 Nov;54:33-50). Shown are the gene symbols (left column) and corresponding protein IDs (right column) of common demyelination-associated proteins.

|  | **Whole protein partners** | **Demyelination-associated protein partners** | **Enrichment factor** | **Adjusted**  **p-value** |
| --- | --- | --- | --- | --- |
| ***TRAF6*** | 0 | 0 | NA | NA |
| ***PRKN*** | 448 | 0 | NA | NA |
| ***HSPA4*** | 363 | 0 | NA | NA |
| ***SQSTM1*** | 294 | 1 | 11.33 | not significant |
| ***HSPA8*** | 708 | 0 | NA | NA |
| ***YWHAZ*** | 412 | 0 | NA | NA |
| ***HSP90AA1*** | 839 | 0 | NA | NA |
| ***YWHAQ*** | 476 | 0 | NA | NA |
| ***CYLD*** | 620 | 0 | NA | NA |
| ***CTNNB1*** | 616 | 0 | NA | NA |
| ***UBC*** | 1050 | 0 | NA | NA |
| ***TP53*** | 1062 | 0 | NA | NA |
| ***CUL7*** | 658 | 0 | NA | NA |
| ***EGFR*** | 1224 | 1 | 2.72 | not significant |
| ***RNF4*** | 1251 | 0 | NA | NA |
| ***MCM2*** | 940 | 1 | 3.54 | not significant |
| ***BRCA1*** | 962 | 0 | NA | NA |
| ***NTRK1*** | 1948 | 1 | 1.50 | not significant |
| ***CUL3*** | 1227 | 0 | NA | NA |
| ***ESR2*** | 2249 | 1 | 1.71 | not significant |

**Legend:** A survey of the human proteome was performed by querying the protein-protein interaction database BioGRID (38). The partners of hub proteins shown to interact with a significantly high number of common neurodegeneration-associated proteins were retrieved. For each of these lists of interactors, a factor of enrichment in common demyelination-associated proteins was established along with an associated Fisher’s exact test p-value, as described in the Materials and Methods section. From left to right, gene symbols and corresponding protein IDs of the identified hub proteins are shown in the first and second column, the associated total numbers of partners and total numbers of common demyelination-associated protein interactors are shown in the third and fourth column. The corresponding enrichment factors and adjusted p-values are shown in the last two columns. Proteins that are detectable by mass spectrometry in human B-cells, according to the “Human Proteome Map” database (41), are highlighted in yellow. NA: not applicable.
